# Natural food intake patterns do not synchronize peripheral clocks

**DOI:** 10.1101/2020.07.28.220509

**Authors:** Xiaobin Xie, Ayaka Kukino, Haley E. Calcagno, Alec Berman, Joseph P. Garner, Matthew P. Butler

**Affiliations:** Oregon Institute of Occupational Health Sciences, Oregon Health & Science University, Portland, OR; Department of Behavioral Neuroscience, Oregon Health & Science University, Portland, OR; Key Laboratory of Protein Modification and Degradation, School of Basic Medical Sciences, Guangzhou Medical University, Guangzhou 511436, China; Department of Comparative Medicine, Stanford University, Stanford, CA

**Author notes:** Corresponding Author: Matthew P. Butler, Oregon Institute of Occupational Health Sciences, Oregon Health & Science University, 3181 SW Sam Jackson Park Road - L606, Portland, OR 97239, |. Author Contributions: Conceptualization, MPB; Methodology, MPB, JPG; Validation, MPB, XX, HG; Investigation, MPB, XX, AK, HG, AB; Writing, MPB, XX; Supervision, MPB.

**Keywords:** meal timing, peripheral clocks, liver, kidney, TRF, time restricted feeding

## Abstract

Food is thought to synchronize circadian clocks in the body, but this is based on time-restricted feeding (TRF) protocols. To test whether naturalistic feeding patterns are sufficient to phase-shift and entrain peripheral tissues, we measured circadian rhythms of the liver, kidney, and submandibular gland in *mPer2*^*Luc*^ mice under different feeding schedules. In ad lib feeding as well as in a schedule designed to mimic the ad lib pattern, PER2::LUC bioluminescence peaked during the night as expected. Surprisingly, shifting the scheduled feeding by 12h caused only small advances (<3h). To isolate the effects of feeding from the light-dark cycle, clock phase was then measured in mice acclimated to scheduled feeding and housed in constant darkness. In these conditions, peripheral clock phases were better predicted by the rest-activity cycle than the food schedule. Under natural feeding patterns, the master pacemaker in the brain sets the phase of peripheral organs independent of feeding behavior.

## Introduction

Healthy physiology is characterized by ∼24h circadian rhythms in all tissues that are entrained (synchronized) to cyclic environmental cues. Disruptions of the circadian system, such as by jet lag and shift work, increase the risk of poor health including obesity, diabetes, and heart disease in both humans and animal models (Karlsson et al., 2001; Turek et al., 2005; Suwazono et al., 2006; Marcheva et al., 2010; Karatsoreos et al., 2011; Pan et al., 2011; Vimalananda et al., 2015). Circadian disruptions cause adverse health outcomes via internal misalignment (in which internal clocks become desynchronized from each other) and external misalignment (in which clocks become desynchronized from external entraining cues) (Davidson et al., 2006; Scheer et al., 2009; Kalsbeek et al., 2011; Karatsoreos et al., 2011; Buxton et al., 2012; LeGates et al., 2012; Leproult et al., 2014; Mukherji et al., 2015b).

Internal clock alignment is maintained by the central pacemaker in the suprachiasmatic nucleus (SCN) of the hypothalamus (Dibner et al., 2010a; Mohawk et al., 2012). The SCN is entrained by light, and in turn, it coordinates the timing of peripheral clocks via several potential pathways, one of which is thought to be the SCN’s control of eating behavior. In studies of time-restricted feeding (TRF), during which rodents are typically exposed to 4-12 h of food availability and 12-20 h of fasting each day, the feeding schedule reliably entrains the circadian phase of peripheral organ clocks, while the SCN remains entrained to the light-dark cycle (Damiola et al., 2000; Stokkan et al., 2001). This has led to a general model of circadian entrainment in which the SCN controls the phase of peripheral clocks via its control of feeding behavior.

TRF imposes long fasting intervals that do not typically occur when food is available ad libitum, and it is not known whether natural eating patterns play a role in entraining peripheral clocks. To test this, we measured peripheral organ rhythms in simulated natural feeding conditions in light-dark cycles and in constant dim light (**Fig. 1, Supplementary Fig. S1**). We focused on the phase of the liver and kidney, two tissues that are entrained by TRF, and the submandibular gland, a tissue with a high amplitude oscillator that is insensitive to TRF and instead entrains to the light-dark cycle (Damiola et al., 2000; Stokkan et al., 2001; Vujovic et al., 2008; Swamy et al., 2018). Surprisingly, we found that simulated natural feeding had little effect on peripheral clocks: peripheral oscillators generally remained entrained to either the light-dark cycle or to the animal’s own rest-activity cycle.

**Figure 1.**
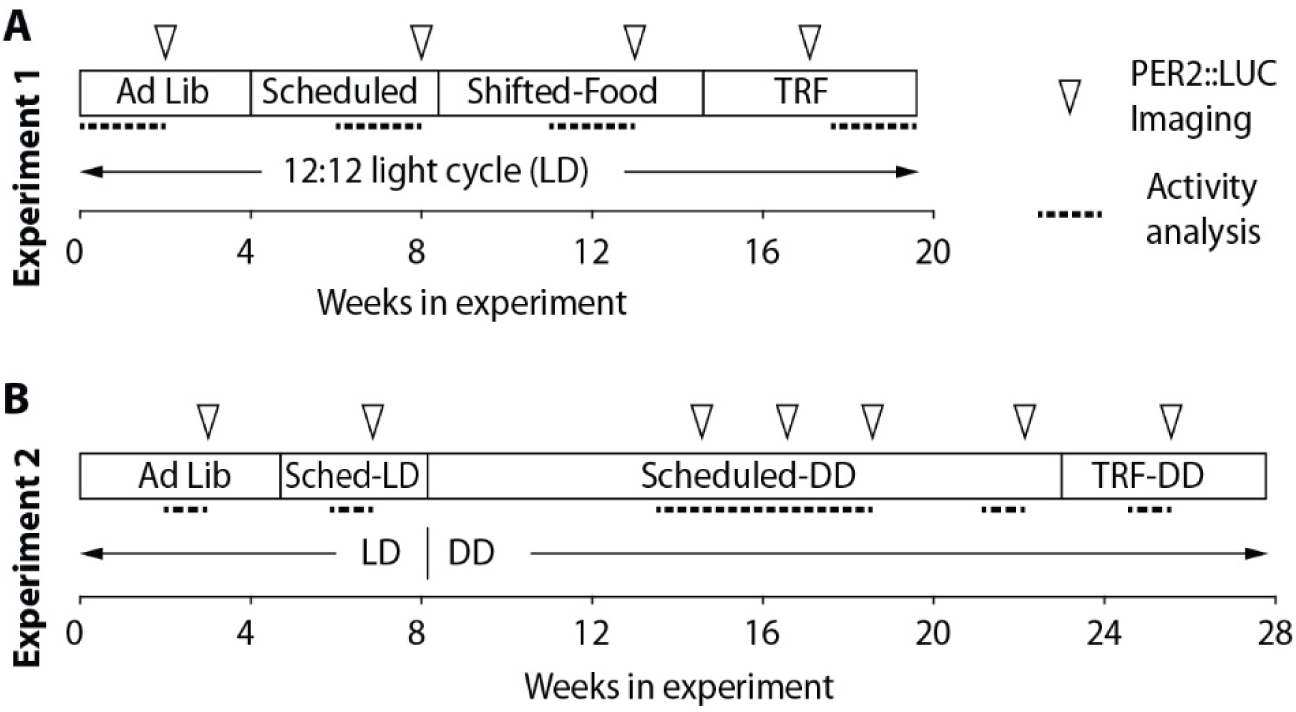
Timeline for both experiments, showing time of baseline intake measures, food provision schedules (Ad Lib, Scheduled, Shifted, or TRF), the light-dark cycle (LD or DD), and points at which PER2::LUC imaging was conducted to measure the phase of peripheral organ circadian rhythms.

## Results

### Experiment 1. Scheduled feeding in light-dark cycles

We first tested whether simulated natural food intake patterns were sufficient to shift locomotor activity or peripheral clocks in a light-dark cycle (LD). After acclimating male mice to custom in-cage automatic feeders (**Supplementary Fig. S2**), the average Ad-Lib intake pattern was calculated. Both meal size and inter-meal interval varied across the light-dark cycle (**Fig 2A-C**), with 65% of food intake during the dark phase. Mean meal size was 10.8 pellets (216 mg) (SD 6.2 pellets) and mean inter-meal interval was 83 min (SD 62). To simulate natural intake patterns, inter-meal intervals were set to 90 min and the amount provided in each meal varied (**Fig. 2D, Supplementary Fig. S3**). Note that because meals were defined by at least 3 pellets, the inter-meal intervals here are longer than the fasting intervals (see Discussion). The group average profile was applied to all mice but the amounts were adjusted to each mouse’s baseline intake. To ensure that mice ate food when it was provided, the total food during Scheduled conditions was reduced by 5% compared to Ad Lib conditions. Body weight was maintained at 99-103% of baseline thereafter (**Supplementary Fig. S4**). The scheduled food was later shifted 12 h (Shifted-Food-LD). Finally, mice were switched to time restricted feeding (TRF-LD), with eight equally sized meals spread across the light portion of the photocycle (**Fig. 2D**).

**Figure 2.**
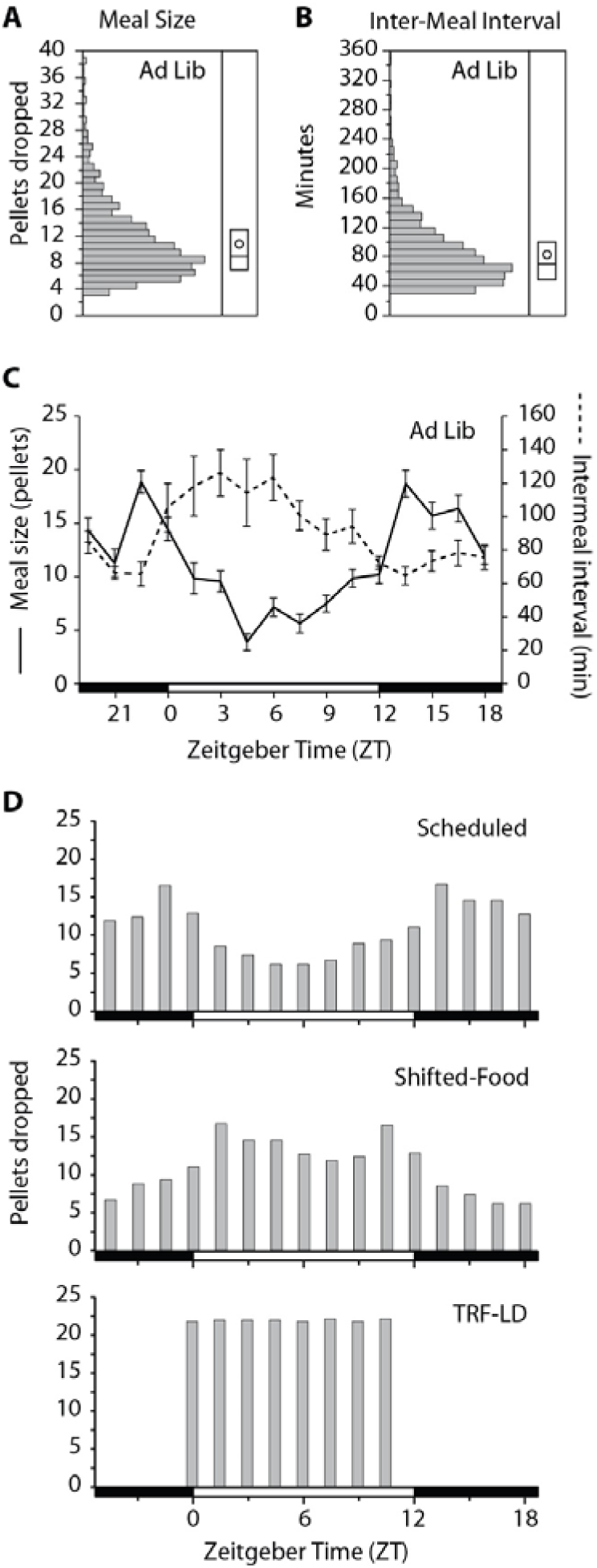
A, B. Distributions of Ad Lib meal sizes in 20 mg pellets and inter-meal interval in minutes (n=10). Box plot shows the median, 25^th^, and 75^th^ percentile. Mean is shown by the circle. C. Both meal size and inter-meal interval varied with time of day during Ad Lib conditions. The light-dark cycle is shown at the bottom, with Zeitgeber Time 0 (ZT0) and ZT12 defined as lights-on and lights-off, respectively. D. The mean pellet drop schedules are plotted for Scheduled, Shifted-Food (12h), and TRF conditions, all conducted in LD.

PER2::LUC bioluminescence rhythms from the liver, kidney, and submandibular gland were assessed in vivo at baseline and during each different feeding schedule (**Fig. 3**). In the group-level analysis, bioluminescence peaked in the late night in all tissues in Ad Lib, Scheduled, and Shifted-Food conditions, suggesting that shifting the food schedule did not phase shift the peripheral clocks.

**Figure 3.**
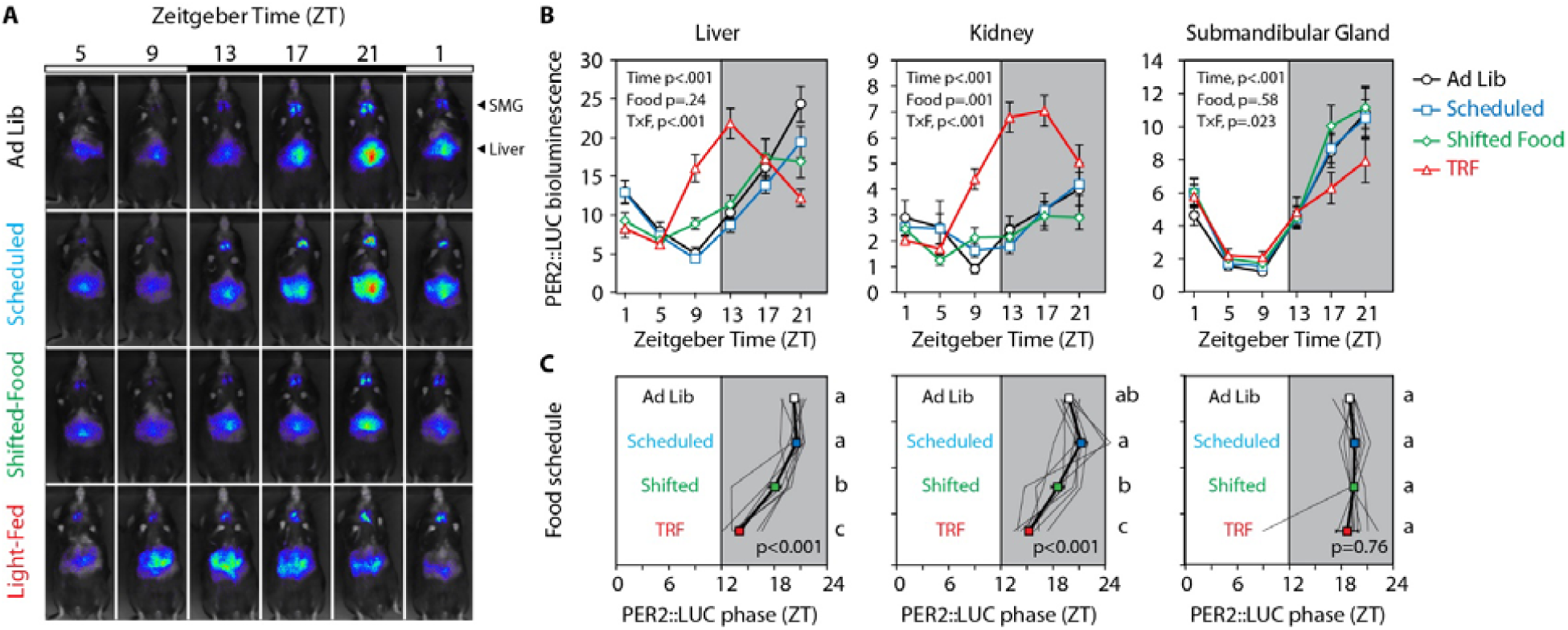
In vivo imaging and determination of activity phase. Every 4 h, mice were imaged after being lightly anesthetized and injected s.c. with luciferin. A. Ventral images from a representative mouse: note that peak PER2::LUC bioluminescence in the SMG occurs at ZT21 in all conditions, whereas peak bioluminescence in the liver occurs earlier only in the TRF light-fed condition. B. Mean bioluminescence is plotted as a function of time of day (n=9). Statistical results from a 2-way repeated measures ANOVA are shown (main and interaction effects of zeitgeber time and food condition). A significant interaction suggests a change in phase or amplitude of the rhythm. C. The phase of each tissue was calculated from the direction of the mean resultant vector and plotted (± SE). P value indicates the result of a 1 way repeated measures ANOVA (feeding condition as independent variable). Conditions that do not share a letter are significantly different (Tukey test, p<.05). Individuals are shown by gray traces.

To capture individual variability, peak phase was calculated for each mouse and tissue (**Supplementary Figs. S5, S6**). In this analysis, there was a significant effect of feeding condition on liver phase (repeated measures ANOVA, F_3,24_ = 44.5, p<.001) and kidney phase (F_3,24_ = 19.6, p<.001), but there was no effect on the submandibular gland (F_3,24_ = 0.39, p=.76) (**Fig. 3C**). Pairwise comparisons showed that Scheduled meals did not change clock phase relative to Ad Lib conditions (Tukey test, n.s.). When the Scheduled feeding was shifted 12 h (Shifted-Food), peripheral tissues continued to peak during the dark, with only small shifts in liver (2.5±0.7 h) and kidney (2.8±1.1 h) (Tukey test, p<.05), and no advance in the submandibular gland (**Fig. 3C**, 0.1±0.4 h, n.s.). The kidneys of one mouse and the liver of another shifted robustly to the new food schedule. Without these, the group phase shifts were less than 2 h. Conclusions based on circular statistics were similar, with the exception that Shifted-Food schedule did not significantly advance the kidney clock (**Supplementary Fig. S5**, Mardia-Watson-Wheeler test, p>.30). These data suggest that naturalistic food intake patterns are not strong enough zeitgebers (time-setting cues) to phase shift the peripheral circadian clock system.

The lack of food entrainment was surprising given previous reports from TRF treated mice (Hatori et al., 2012; Tahara et al., 2012; Swamy et al., 2018), so mice were switched to a 12h food availability cycle, with 8 equally sized meals presented during the 12 hours of the light cycle (ZT0-12). After 2 weeks, this schedule was sufficient to significantly advance the liver and kidney clocks relative to the first Scheduled condition by 6.5 and 6.0 h (**Fig. 3C, Supplementary Fig. S5**). TRF had no effect on submandibular gland phase.

During different food conditions, mice remained nocturnal as expected (Hatori et al., 2012; Bray et al., 2013). The nocturnality index (nocturnal to diurnal activity ratio) was significantly greater than 1 in all conditions, though there was a significant reduction in this index during the last TRF-LD condition (**Supplementary Figs. S7, S8**).

### Experiment 2. Scheduled feeding in constant darkness

In Experiment 1, shifting the simulated natural food intake did not appreciably shift locomotor activity or peripheral tissue clock phase. But the zeitgeber strength necessary to shift clocks may be much higher than to maintain entrainment, for example as observed in responses to light (Butler and Silver, 2011). Therefore, Experiment 2 was conducted on a new cohort of mice to determine if simulated natural intake patterns were sufficient to entrain the liver clock in the absence of a light-dark cycle.

Mice exhibited a normal feeding pattern with 68% of food consumed during the dark phase (**Fig. 4**). The average intake profile for the second cohort of mice was again applied using a 90 min inter-meal interval. Body weight remained stable (**Fig. S9**), and transitioning from Ad Lib to Scheduled feeding in LD did not phase shift the circadian system (**Fig. S10**).

**Figure 4.**
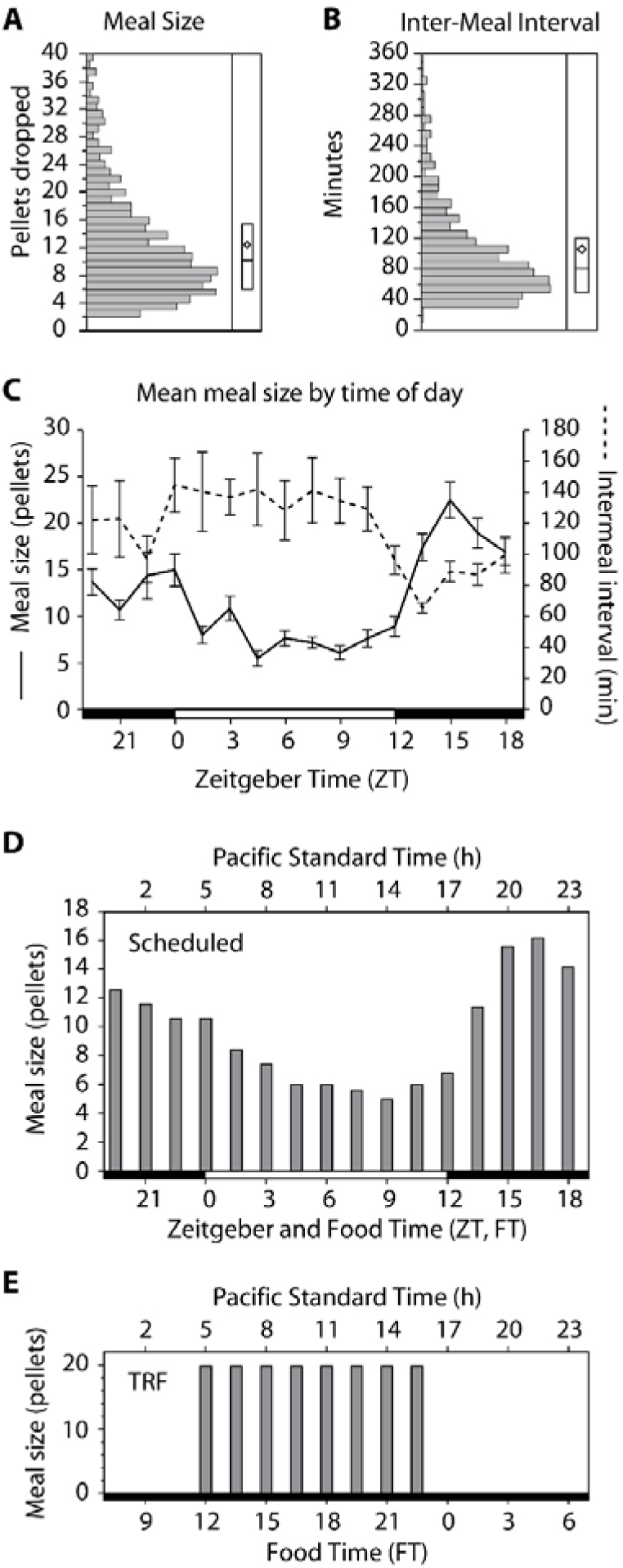
A, B. Experiment 2 meal size and inter-meal interval distributions as in Fig. 2. C. Both meal size and inter-meal interval varied with time of day during ad lib conditions. D. The mean food rate was calculated for n=12 mice and applied to all mice, with mean level adjusted for individual intake at baseline. The peak in intake at the end of the night was not as pronounced as in Experiment 1. Here ZT12 and FT12 occur at 1700 PST. E. After free-running in darkness with scheduled feeding, mice were switched to TRF conditions with 12 h food availability with FT12 at 0500 PST.

Without changing the Scheduled food condition, the mice were released into constant dim red light (DD, 0.1 lux), and their activity rhythms began to free-run according to their endogenous non-24h period. This immediately established that simulated natural food patterns do not entrain locomotor activity rhythms. In DD, activity phase and peripheral organ phase were calculated periodically over 3 months, and the PER2::LUC bioluminescence was analyzed as a function of the rest-activity cycle (circadian time, with activity onset defined as CT12) or food time (FT12-24 representing the period of greatest intake/provision and corresponding to the original 1700-0500 dark period) (**Fig. 5**). PER2::LUC expression was synchronized to the rest-activity cycle: circadian time explained 36-70% of the variance in the model (**Supplementary Table S1**). In contrast, peripheral rhythms were desynchronized when plotted against the food intake cycle, and food time only explained 2-9% of variance.

**Figure 5.**
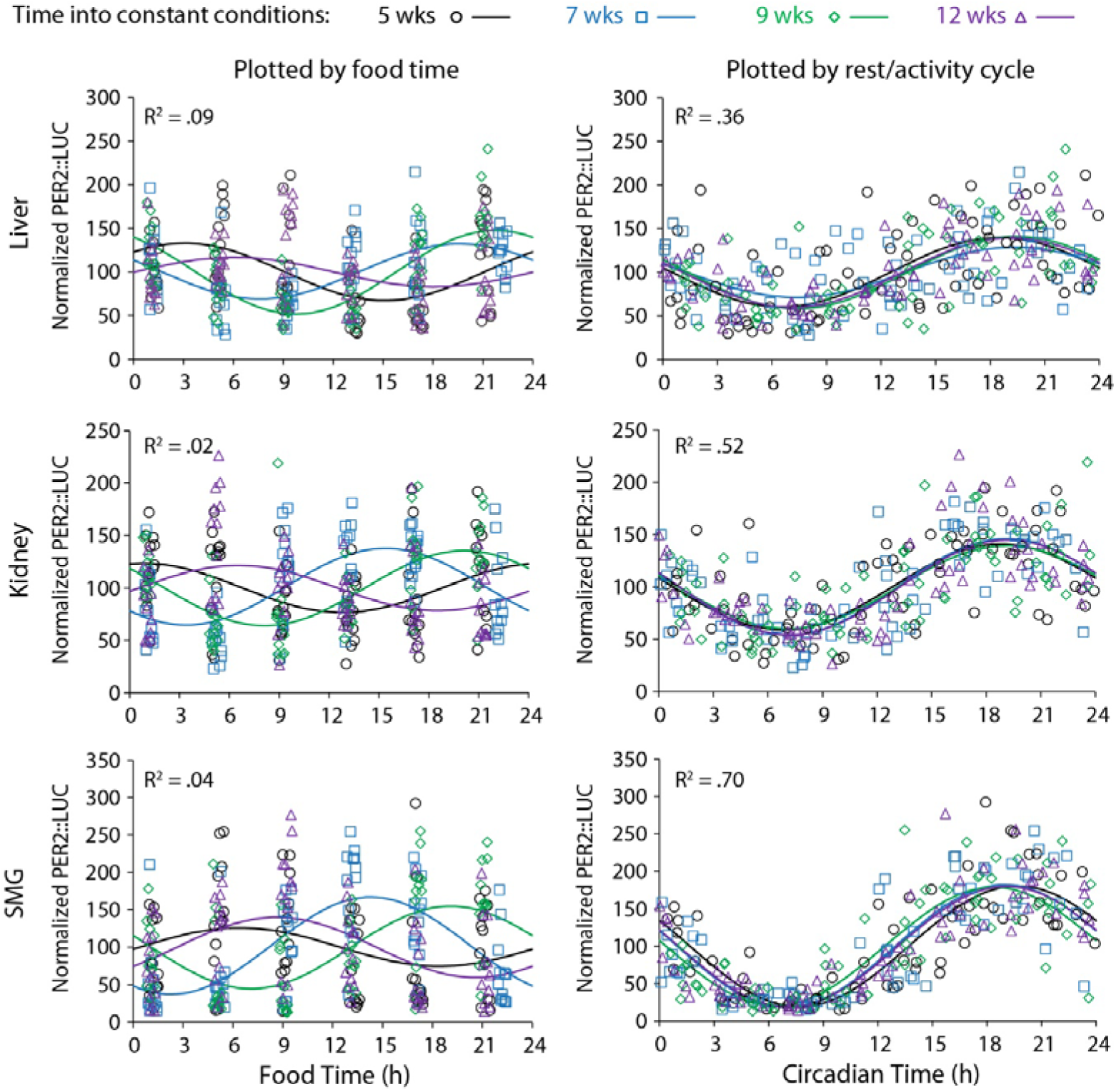
PER2::LUC bioluminescence was measured *in vivo* 6 times/day (n=12 mice). This was done after 5 (circle), 7 (box), 9 (diamond), and 12 weeks (triangle) of housing in constant conditions. The same data are plotted against either the timing of food (left) or the timing of each animal’s rest-activity cycle (right). Curves are the best fit cosines for each measurement point. R^2^ values are from the full model that combined all four measurement days (**Supplementary Table S1**). The liver and kidneys are much better aligned with circadian rhythms of rest/activity, and thus with the brain’s pacemaker, than with the schedule of food intake.

Over the four tests of phase during Scheduled-DD conditions, each tissue’s phase was calculated and plotted circularly as a function of food time or circadian time (**Supplementary Fig. S11**). Scheduled food did not entrain peripheral organs. Instead, from test to test, the phase in food time varied widely (Mardia-Watson-Wheeler test, W≥14.9, p≤.021). In contrast, peripheral tissues were always in phase with circadian time with no test-to-test variability (W≤10.7, p≥.099).

There was evidence for individual variability in sensitivity to food cues (**Supplementary Fig. S12**). For each tissue, we tested whether circadian phase clustered better along food time or circadian time by calculating the overall mean resultant vector from the four phases measured during constant conditions (4 repeated measures of phase per tissue). Submandibular glands did not entrain to food and always showed significant clustering in circadian time (**Fig. 6**, r≥.83, p<.05 for all). In contrast, phase in some kidneys and livers was sensitive to food intake patterns. One kidney and one liver (not from the same mouse) showed significant phase clustering in food time, indicating food entrainment. Nevertheless, 8/12 livers and 11/12 kidneys showed stronger clustering by circadian time (above the dotted line in **Fig. 6**); this was significant in 6/12 livers and 9/12 kidneys.

**Figure 6.**
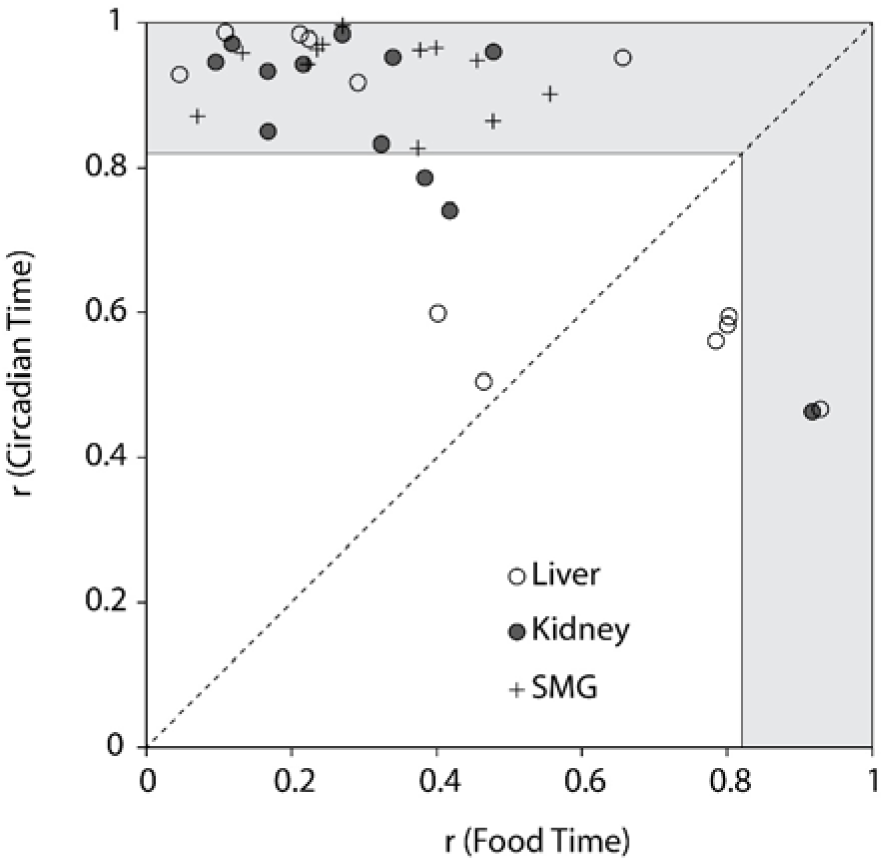
Individual variability in liver and kidney entrainment to food. For each tissue in each mouse, a preferred phase was calculated and r calculated from the four measures completed in constant conditions. This was done both in food time and in circadian time. The two r statistics are plotted against each other. Points above the diagonal dotted line indicate that circadian time better predicts phase than food time. Points that fall in the grey shaded area have a significant preferred phase (r>0.82; p<0.05; Rayleigh test for n=4). Note: the liver and kidney with significant phase clustering by food time are not from the same mouse. Activity and tissue phases for the mouse with significant liver food time (empty circle on right) are shown in **Supplementary Fig. S12A**.

After 15 weeks in Scheduled-DD conditions, mice were switched to TRF with 12 h of feeding and 12 h of fasting, to determine whether restricted feeding with a long daily fast could entrain rhythms in DD. As in Experiment 1, mice were provided 8 equally sized meals presented during the time of the original photophase (0500-1700 PST). Period shortened in the majority of mice, but locomotor activity entrained (period = 24h) in only two mice (**Supplementary Fig. S13**).

Though food schedules did not entrain locomotor activity in most mice, TRF did entrain rhythms of the liver and kidney (**Supplementary Fig. S14**). When tissue phases were plotted by food time, the liver and kidney phases strongly clustered around FT0 (Rayleigh test, p<.001), similar to phases observed during Ad Lib conditions. Though the SMG displayed statistically significant clustering, there was a wide range of phases. The opposite results were obtained when plotted by circadian time: strong clustering for the SMG and weak clustering for the liver and kidney. Therefore, TRF in DD effectively entrained kidney and liver but did not entrain either the SMG or locomotor activity.

## Discussion

We show here that natural feeding behavior has little resetting effect on peripheral clocks. In Experiment 1, a 12-h shift of the simulated natural food provision schedule resulted in small shifts in liver and kidney and no shift in the SMG. In Experiment 2, we removed the light-dark cycle and tested whether the scheduled food would continue to entrain peripheral clocks. Peripheral organ phase was nevertheless much better predicted by the rest-activity cycle; food significantly entrained only 1 of 12 kidneys and 1 of 12 livers. As a positive control, TRF was an effective zeitgeber in both experiments, unlike the simulated natural feeding. From these data, we infer that under normal conditions of food intake, the SCN remains the dominant synchronizer of peripheral clocks and that this does not require the intermediate step of feeding behavior.

In TRF experiments, food reliably entrains peripheral organs (Damiola et al., 2000; Stokkan et al., 2001). This has led to a model of peripheral entrainment in which the SCN entrains patterns of behavior (activity and food intake), and these behaviors and their metabolic sequelae in turn entrain peripheral organs. In this experiment of simulated natural feeding, however, the effects of food intake rhythms on peripheral phase were small. This may be because natural intake patterns of rodents do not feature long fasting intervals. The extended fast may be necessary to entrain peripheral clocks via reduced circulating insulin (Mukherji et al., 2015a). The long fasts are also important for the metabolic health benefits of TRF in rodents and humans (Hatori et al., 2012; Chaix et al., 2014; Gill and Panda, 2015; Paoli et al., 2019; Stenvers et al., 2019).

Fasting intervals were longer during the day than the night, so by setting a constant inter-meal interval (90 min), we may have removed a physiologically relevant fasting signal. The shortest fast needed to entrain peripheral clocks is not known, but is likely between 4 and 8 h, given that 8h but not 4h fasts alter liver clock gene expression (Mukherji et al., 2015a). We therefore analyzed the distribution of fasting intervals during Ad Lib conditions to determine how often >4h fasts occur. True fasting intervals (no pellets dropped) are shorter than the inter-meal intervals, since mice occasionally consumed 1-2 pellets and this was not counted as a meal. The maximum average fasting intervals per bin in the two experiments were 93 min (ZT4.5-ZT6) and 91 min (ZT3-4.5), similar to the 90 min inter-meal interval imposed on the mice during scheduled feeding and shorter than the peak mean inter-meal intervals (120–140 min, **Figs. 2, 4**). Of course, the averages may not be informative if individual mice still have long fasts each day. The incidence of long fasts are shown in **Supplementary Tables S2-S3**. Over 80 mouse-days under Ad Lib conditions in Experiment 1, only 13 intervals lasted longer than 4 hours (0.7% of all fasting intervals). In Experiment 2, there were 53 such fasts in 120 mouse days (2.3%). Based on TRF experiments, it would be the long fasts during the light phase that would entrain normal night-time peaks of PER2::LUC in peripheral organs. These were not equally distributed across mice. Though two mice in Experiment 2 had almost one long fast per day, the other 20 mice had long fasts on half of their days or fewer and five never experienced a light-phase fast >4 hours. These data match other reports that voluntary inter-meal intervals in mice rarely exceed 5 h (Jensen et al., 2013). We therefore conclude that naturally occurring fasting intervals during Ad Lib feeding are unlikely to entrain peripheral clocks.

Our results complement investigations of meal frequency effects on clocks. Kuroda et al. (2012) show that both meal frequency and size affect the phase angle of entrainment in the liver, kidney, and submandibular gland. Phase nevertheless generally tracked with the longest fast each day (8-12h). In experiments of equally spaced meals to eliminate the rhythmic food signal, peripheral rhythms remain entrained to the SCN in rats, mostly likely via adrenal hormone signaling (Kalsbeek and Strubbe, 1998; Su et al., 2016). In mice, peripheral clock gene rhythms are preserved but advanced ∼2-4 h under the 6 meal/day schedule compared to ad libitum feeding (Kuroda et al., 2012; Sen et al., 2017); the amount of advance seems to increase with caloric restriction. These data establish that feeding rhythms are not necessary to entrain peripheral clocks, but do not speak to the relative importance of meal-induced cues versus other SCN-derived cues in ad lib fed mice. In turn, our results suggest that even in the presence of naturally occurring food-derived metabolic signals, more proximate SCN-controlled cues orchestrate the peripheral clock system. The SCN communicates time of day information via neural, endocrine, and physiological pathways (Dibner et al., 2010b; Mohawk et al., 2012). Potential entrainment mechanisms include glucocorticoids (Balsalobre et al., 2000; Valenzuela et al., 2008; Pezuk et al., 2012; Cuesta et al., 2015), autonomic nervous system projections (Ueyama et al., 1999; Cailotto et al., 2005), body temperature (Brown et al., 2002; Buhr et al., 2010), and melatonin (von Gall et al., 2005).

Two results from the TRF conditions bear discussion: peripheral phase in Expt. 1 and the differential entraining effect on clocks in Expt. 2. In Experiment. 1, the liver and kidney shifted ∼6h, substantially less than the ∼12h expected based on restricted feeding (Damiola et al., 2000; Stokkan et al., 2001; Vollmers et al., 2009) and which we have observed when mice can eat ad libitum during food availability periods (Swamy et al., 2018). A key difference may be the use of equal-sized and -spaced meals: similar 4-7h shifts were reported in mice fed 3 meals in either the light or dark {Kuroda, 2012 #2348}. Set meal sizes prevent the initial gorging response to food availability that is only observed in light-fed mice (Bray et al., 2013). These data suggest that both long fasts and a large refeeding response are necessary to fully reset clocks in day-fed mice. In Experiment 2, locomotor activity rhythms continued to free-run in a majority of mice despite the 12h feeding schedule during TRF-DD. This is consistent with some reports (providing 2-6 h of food availability per day) (Abe et al., 1989; Mendoza et al., 2010; Takasu et al., 2012) and differs from others (4-8 h of food/day) (Feillet et al., 2006; Sheward et al., 2007). Others have shown an intermediate proportion of mice that entrain to food (44-67%) (Marchant and Mistlberger, 1997; Refinetti, 2015). Both strain differences and duration of daily food availability can contribute to this inconsistency (Castillo et al., 2004). The misalignment between liver/kidney (entrained to food) and activity/submandibular gland (free-running and synchronized to each other) further underscores that TRF has little effect on the central clock.

To what extent do our results shed light on human peripheral clock control? It is not yet known whether meals entrain peripheral clocks in humans as they do in mice during TRF because there are few methods to directly measure peripheral clock phase in humans. Studies with shifted meal times have demonstrated that meals set the phase of glucose and other metabolite rhythms, but the effects on clock phase in white blood cells is much smaller (Wehrens et al., 2017; Skene et al., 2018) – indeed, clock gene phase shifts are similar to what we observed in mice with physiologically relevant feeding pattern (Experiment 1). Though blood markers may reflect an averaging of phase information from many tissues, the results suggest that meals are weak zeitgebers for human clocks even as they drive daily rhythms in metabolic physiology. The difference may lie in how mice and humans respond to fasting. Unlike in humans, an overnight fast in a mouse can reduce body weight by 15%, induce a catabolic state, and engage starvation protection mechanisms (Ayala et al., 2010; Jensen et al., 2013). Therefore, even though TRF is a better match for human food intake patterns, ad lib-fed mice may better mimic the gentler waxing and waning of circulating metabolic markers over the day in humans. It will be of interest then to determine whether the SCN’s control of peripheral rhythms via non-food pathways is stronger in humans than in mice.

This experiment was enabled by home cage automated feeders that allowed precise measurement of food intake and provision of a set pattern. The use of *mPer2*^*Luc*^ mice and in vivo imaging let us repeatedly sample circadian phase in multiple tissues in individuals, revealing some inter-individual variability in zeitgeber sensitivity. Glucose, insulin, and other potential resetting cues were not measured in this study, so although simulated natural food patterns did not alter behavior and peripheral organ rhythms compared to ad lib intake, whether and how the food schedule may have altered metabolic signaling remains unknown. Finally, the scheduled food pattern was imposed by varying meal size, so the contribution of inter-meal interval variability remains unknown.

These two experiments show that natural food intake patterns have little effect on the phase of peripheral clocks. These data challenge a major model of circadian clocks in which peripheral clocks are coordinated by the SCN via its control of food intake. Instead, our results suggest that during normal feeding conditions, the SCN entrains peripheral clocks via feeding-independent pathways.

## Methods

### Animals and housing

*mPer2*^*Luc*^ mice on a C57Bl/6 background, bearing a knockin fusion protein combining the clock gene *Period2* and a firefly luciferase reporter, were purchased from Jackson Laboratories (B6.129S6-*Per2*^*tm1Jt*^/J, Strain Code: 006852) (Yoo et al., 2004). Male mice for this experiment were bred locally and maintained in a Thoren ventilated caging system in a pathogen-free barrier facility on pelleted cellulose bedding (BioFresh Performance Bedding, ¼” pelleted cellulose, Absorption Corp, Ferndale, WA), with food (LabDiet 5L0D) and water available *ad libitum.* For experiments, adult mice were transferred to an animal enclosure outside of the barrier facility, and housed individually in cages with a custom pellet feeder (Telos Discovery Systems, West Lafayette, IN, **Supplementary Fig. S2**), in a single cabinet with light-dark cycle control (Phenome Technologies, Lincolnshire, IL). The home cage feeder stands at one end of the cage (Thoren model #1, 19.6 cm×30.9 cm×13.3 cm), and takes up 7.5 cm, therefore limiting the living space length to 23.4 cm. Flat wire lids were used above the cage, and to prevent the water bottle from interfering with activity recording (see below), the water bottle was mounted at the end of cage, with the sipper tube extending through the watering grommet (7.5 cm from the cage floor). Light was provided by green LEDs (525nm, full width at half maximum (FWHM) 25nm, 125 lux) from 0500-1700 PST. To aid in husbandry and to ameliorate potential dim-light induced effects on the circadian system during imaging procedures, a dim red light was on continually in the cabinet (625nm, FWHM 25nm, 0.2 lux). All procedures were approved by the Institutional Animal Care and Use Committee of Oregon Health & Science University.

### Feeding

At specified times, the feeders automatically dispensed 20 mg food pellets (Bio Serv. Rodent Grain-Based Diet Sterile Product #S0163). The presence of the pellet and head entries into the pellet trough (1.5cm × 1.5cm) were recorded by infrared beam breaks. During ad libitum feeding, a new pellet was dropped each time the pellet in the trough is removed. A failsafe mechanism drops another pellet 10 sec later if the first pellet is not detected in the trough. Each dispensing event, head entry, and pellet removal is time stamped and logged by an Access 2010 database (Microsoft, Seattle, WA); analyses were then conducted on binned counts (10 min bins). Individual meals were identified in the record by feeding bouts of at least three pellets (≥60 mg) with at least 20 min of fasting preceding and following the meal. Feeders were checked periodically to ensure that all pellets were eaten prior to the next meal and that no animals were hoarding food. Troughs were empty in 92% of 278 observations suggesting that the dispensing schedule determined the intake schedule.

### Locomotor Activity

Activity counts were measured by a passive infrared detector (Phidgets 1111_0 motion sensor) mounted 16.5 cm above the cage; data were collected as counts per 10 min bin. Activity was used to calculate the activity profile (mean activity per hour over 14 days) and nocturnality index (dark phase activity divided by light phase activity over 14 days). Additionally, circadian period and phase during constant conditions were calculated from consecutive activity onsets over 7-10 days in ClockLab. Activity onset was defined as Circadian Time 12 (CT12).

### Protocol schedules

<u>Experiment 1</u> The timeline of events is shown in **Fig. 1A**. At week 0, mice (21-23 weeks of age, n=10) were transferred to the automatic feeding cages. Mice were maintained in ad libitum (**Ad Lib**) feeding conditions with a 12:12 light:dark (**LD**) schedule for 4 weeks. From weeks 4-9, mice were exposed to Scheduled Feeding (**Scheduled-LD**) that simulated the group’s average feeding profile. At week 9, the scheduled feeding pattern was shifted by 12 h (**Shifted-Food-LD**). Finally, at week 15, mice were exposed to time-restricted feeding (**TRF-LD**) for 5 weeks, in which food was provided only during lights-on. Peripheral PER2::LUC bioluminescence rhythms were measured once in each feeding condition. For one mouse, a feeder failure introduced long fasts during the Scheduled and Shifted-Food conditions. The mouse recovered but his data were excluded a priori from peripheral clock phase analyses, leaving an analytic dataset of n=9. <u>Experiment 2.</u> Mice, aged 16-17 weeks (n=12), were transferred to automatic feeding cages as above, and fed **Ad Lib** for 5 weeks (**Fig. 1B**). Scheduled feeding was imposed at week 5, simulating the ad lib intake pattern for this new cohort (**Scheduled-LD**). From week 8, the mice were housed in constant dim red light (**Scheduled-DD**, 0.1 lux). Finally, they were shifted to TRF in DD (**TRF-DD**); food was available during the initial time of lights-on (0500-1700 PST). As above, peripheral organ rhythms were measured from PER2::LUC bioluminescence.

### Imaging protocol

An in vivo imaging system (Stanford Photonics, Stanford, CA) was used to measure PER2::LUC bioluminescence in the liver, submandibular gland, and kidney (Tahara et al., 2012; Swamy et al., 2018); this system includes computer assisted capture of bioluminescent images via an Electron Magnified (EM) CCD camera (ImageEM, Hamamatsu, Japan, controlled by Piper software version 2.6.89.18, Stanford Photonics) connected to an ONYX dark box (Stanford Photonics) in which mice can be maintained under isoflurane anesthesia on a 37°C temperature controlled stage (mTCII micro-Temperature Controller, Cell MicroControls, Norfolk, VA). Clock phase was determined from PER2::LUC bioluminescence measured every 4 h (6 measures in one day). Mice were anesthetized with 2-4% isoflurane to prevent movement during imaging. Prior to taking the first image, the fur was shaved from surfaces close to the liver, submandibular gland, and kidneys. Mice were then injected s.c. (15mg/kg) with D-luciferin potassium salt (Promega, Madison, WI) dissolved in sterile phosphate buffered saline (30mg/10mL) and filtered (0.2µm). The dorsal and ventral surfaces of the mice were imaged 8 and 10 min after luciferin injection, respectively. Bioluminescence was captured by the camera in EM mode (sum of eight 125 ms exposures, gain 500). A brightfield reference image was taken each time under dim red light (633 nm, FWHM 15 nm, 1.6 lux). After imaging, the animals were immediately returned to their cages for recovery.

Bioluminescence was scored offline. Each image was opened in ImageJ (NIH), and the intensity of the 24 bit greyscale image quantified using defined regions of interest, centered on the brightest areas of the tissue (liver: 6 mm × 6 mm, both kidneys: 25 mm × 20 mm, submandibular gland: 10 mm × 10 mm).

### Statistics

Error is reported as SEM unless otherwise indicated, and all statistical tests were evaluated as two-sided. <u>Defining time.</u> This experiment combined light cycles, food cycles, and free-running cycles, so data were analyzed as a function of zeitgeber time, food time, and circadian time, respectively (**Supplementary Fig. S1**). Zeitgeber time (ZT) is defined by the light-dark cycle with lights-off at ZT12 and lights-on at ZT0. Food time (FT) is defined similarly, with the major intake occurring from FT12 to FT0. For example, in Scheduled-LD conditions, FT12 = ZT12. During TRF, FT12 is defined as the start of food availability. Lastly, Circadian Time (CT) is specific to each animal’s rest/activity cycle, and CT12 is defined as the time of activity onset. <u>Assessing rhythms</u>. The six in vivo bioluminescence measurements per 24 h were used to calculated a circadian profile and a peak phase. Because of variation in skin pigmentation and fur shaving, the six measures were normalized as percent of mean within mouse within organ within experimental time point (**Supplementary Figure S6**). *Circadian profile comparisons*: In Experiment 1, circadian profiles of bioluminescence were analyzed by repeated measures ANOVA with food condition and zeitgeber time as independent factors. In Experiment 2, bioluminescence patterns were analyzed by cosinor analysis as a function of circadian time or food time. A mouse’s activity rhythm free-runs in constant conditions, so circadian time drifted in and out of phase with the 24 h scheduled food time. The ability of circadian time versus food time to predict peripheral organ phase was assessed by comparing goodness of fit of the cosinor regression (R^2^ and AICc). *Phase analysis*: Circadian phase of individual organs was defined by the direction of the mean resultant vector (**Supplementary Fig. S6**); this provided point estimates of phase for each tissue in each mouse. Circular statistics were employed to determine whether there was a preferred phase in each condition (Rayleigh test) and whether tissue phase differed between feeding conditions or between different weeks into DD (Mardia-Watson-Wheeler test) (Batschelet, 1981). <u>Individual differences.</u> To analyze individual differences, phase clustering within mouse and within organ was assessed by the Rayleigh test (n=4 phase determinations per mouse per organ in DD). The mean resultant vector lengths (r: r=1 when all points occur at the same phase, r=0 when points are uniformly distributed around the clock) were plotted to illustrate the strength of phase clustering when assessed by circadian time versus food time during DD (e.g., **Fig. 6**). <u>Other behaviors</u>. Nocturnality was analyzed by ordinary regression on log-transformed activity with mouse as a repeated measure, and are presented as median and 95% confidence interval. Groups are more nocturnal if the 95% confidence interval does not include 1. Changes in circadian period and phase were assessed by paired t-test.

## Data availability

The datasets generated and analyzed during the current study are available from the corresponding author on reasonable request.

## Acknowledgements

The authors thank Drs. Charles N. Allen, Steven A. Shea, Debra J. Skene, Andrew W. McHill, Mitchell S. Turker, Thijs J. Walbeek, and Brady Kalb for helpful discussions over the course of this work. This work was supported in part by grant NS102962, a New Investigator Grant from the Medical Research Foundation, and by the Oregon Institute of Occupational Health Sciences via funds from the State of Oregon (ORS 656.630).

