## Supplementary for "Natural food intake patterns do not synchronize peripheral clocks" (from another version of this preprint: version 1, 2020-07-29)

**Supplementary Figure S1.** Defining Time. This schematic shows the relationship of different timing conventions for each stage of the experiments. Zeitgeber Time (ZT) indicates the light-dark cycle, with ZT12 at lights-off. Food Time (FT) indicates the food schedule with FT12 corresponding to the start of the primary food intake period or the start of food availability during time-restricted feeding (TRF). Circadian Time (CT) indicates the rest-activity cycle in constant conditions, with CT12 at locomotor activity onset. A. In Experiment 1, Food Time shifts but Zeitgeber Time does not. B. In Experiment 2, Food Time remains constant until shifted for the TRF-DD conditions. When mice are free-running, the rest-activity cycles, and therefore Circadian Time, drift with respect to Food Time and external clock time (PST). Dark gray rectangles illustrate active periods of the mice.

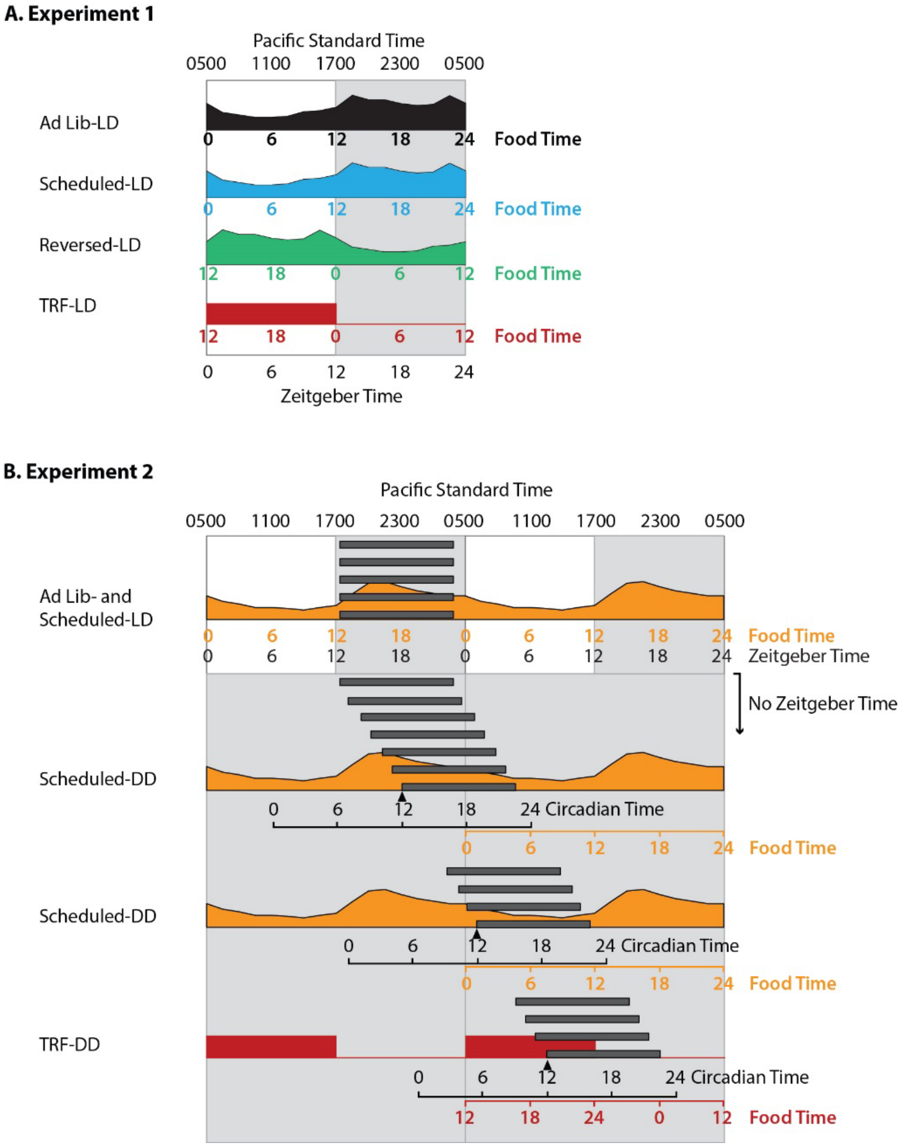

**Supplementary Figure S2.** Automatic feeders were designed to fit in the mouse home cage. Food pellets (20 mg) are stored in hoppers on the back and are dropped one at a time into a food trough. One infrared beam detects head entries into the trough; the second senses the presence or absence of a pellet. The system also records pellet dispensing, humidity, temperature, light, and general locomotor activity.  B. 72 h of representative data from a mouse under Ad Lib feeding conditions. Lights turn on and off at Zeitgeber Time (ZT) 0 and 12 respectively.

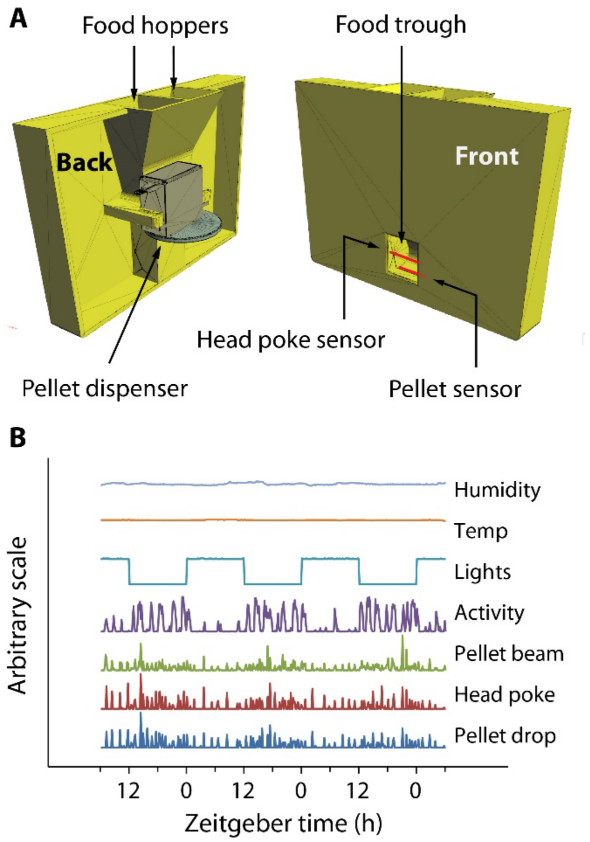

**Supplementary Figure S3.** Both meal size and inter-meal interval varied in Experiment 1 (see **Fig. 2A-C**). In selecting a simulated natural feeding schedule, we examined inter-mouse variability to determine which was more stable. The raw data for the 10 mice contributing to manuscript **Fig. 2C** (below, top) suggested much more inter-mouse variability in the inter-meal interval than in meal size. To quantify the similarity of the curves, we calculated cosine regressions to the raw data for each mouse (middle), and then rescaled these (bottom, translation around the origin and uniform scaling along the y axis). Following this, the root mean square error (RMSE) across meal size curves is half that of the inter-meal interval curves, suggesting that variation in meal size is a more uniform daily signal within the cohort. We therefore concluded that meal size is more likely to be a robust physiological signal in the context of daily resetting.

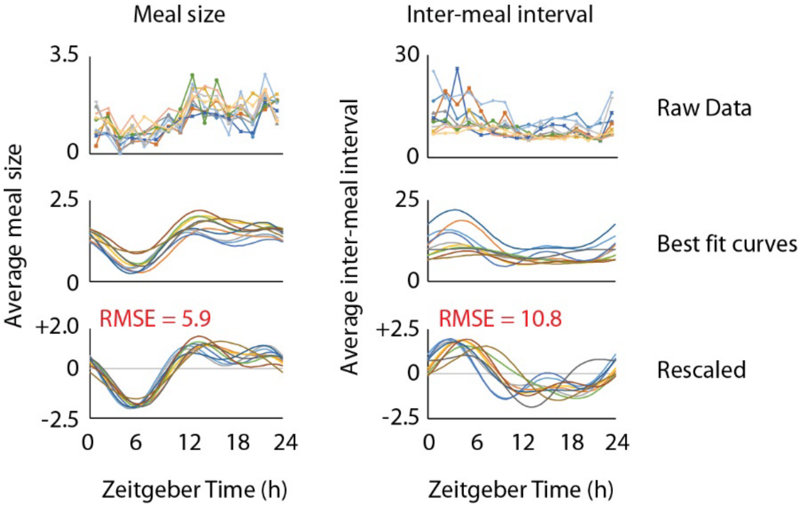


**Supplementary Figure S4.** In Experiment 1, mean body weight (± SE) remained stable. Some variability is due to differences in the time of day of weighing.

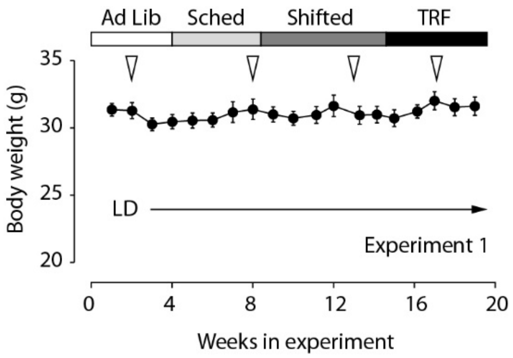

**Supplementary Figure S5.** Left. Pseudocolored bioluminescence from the liver, right kidney, and SMG for the mouse shown in Fig. 3. Scale bar, 5mm. Right. Each tissue's phase is plotted around the circumference of the circles relative to the light-dark cycle. The mean phase within condition is shown by the mean resultant vector. Phase differences between conditions were tested by the Mardia-Watson-Wheeler test, and significance indicated: *p<.05, **p<.01, ***p<.001. Below: Results of the Mardia-Watson-Wheeler test for differences in the phase distribution across food conditions are shown for the overall 4-condition comparison, and for four planned pairwise comparisons. Shifting the food 12 h (blue to green vector) caused a small advance in the liver but no significant advance in the other tissues.

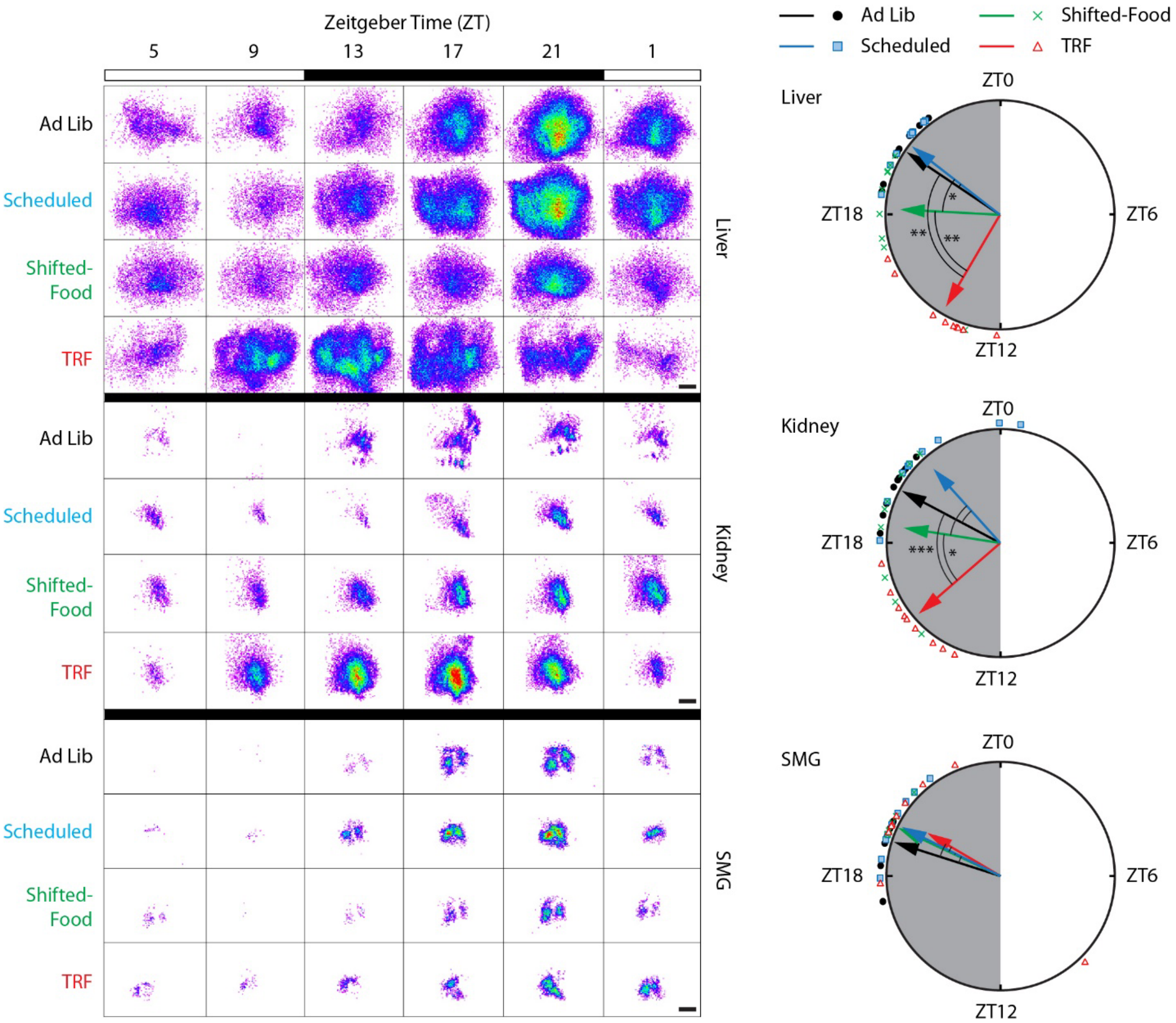


Liver

Overall: W = 29.9, p < .001 ***

Ad Lib | Scheduled: $\chi^2 = 1.1$, p = .57

Scheduled | Shifted: $\chi^2 = 6.7$, p = .034 *

Shifted | TRF: $\chi^2 = 10.8$, p = .004 **

TRF | Ad Lib: $\chi^2 = 12.3$, p = .002 **

Kidney

Overall: W = 24.5, p < .001 ***

Ad Lib | Scheduled: $\chi^2 = 4.6$, p = .10

Scheduled | Shifted: $\chi^2 = 2.4$, p = .30

Shifted | TRF: $\chi^2 = 6.2$, p = .046 *

TRF | Ad Lib: $\chi^2 = 13.9$, p < .001 ***

SMG

Overall: W = 6.6, p = .36

Ad Lib | Scheduled: $\chi^2 = 1.5$, p = .48

Scheduled | Shifted: $\chi^2 = 4.6$, p = .10

Shifted | TRF: $\chi^2 = 2.6$, p = .27

TRF | Ad Lib: $\chi^2 = 1.6$, p = .44

**Supplementary Figure S6.** Example of phase determination via the mean resultant vector. Raw bioluminescence images are measured and converted to normalized units (A, %mean). Normalized data are re-plotted in a circular coordinate system, and the mean resultant vector is calculated. The direction of the vector is the mean phase (ZT 18.98 for this liver example).

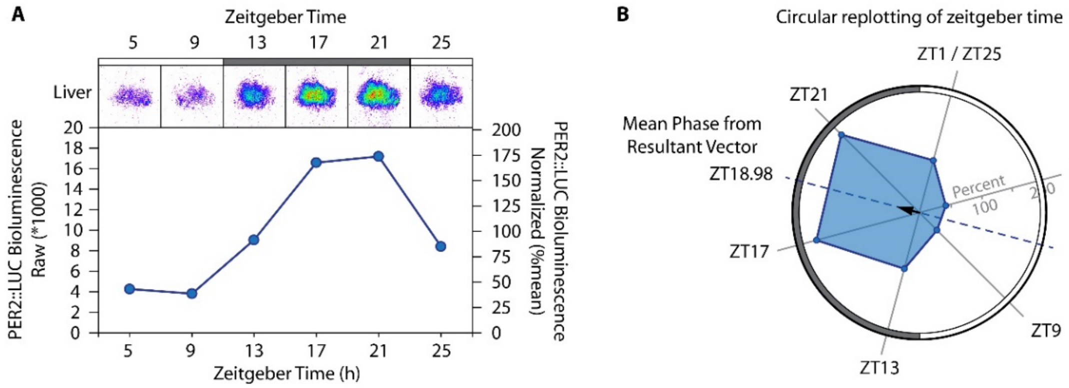


**Supplementary Figure S7.** Representative general locomotor activity, pellet drops, and head pokes for one mouse during Experiment 1. During all feeding conditions, the mouse remained nocturnal. Vertical hatch marks on each line show activity, pellet drops by the feeder, or head pokes by the mouse into the food trough. Consecutive days are aligned from top to bottom. The data are double-plotted for better visualization. The black and white bars at the top show the light-dark cycle.

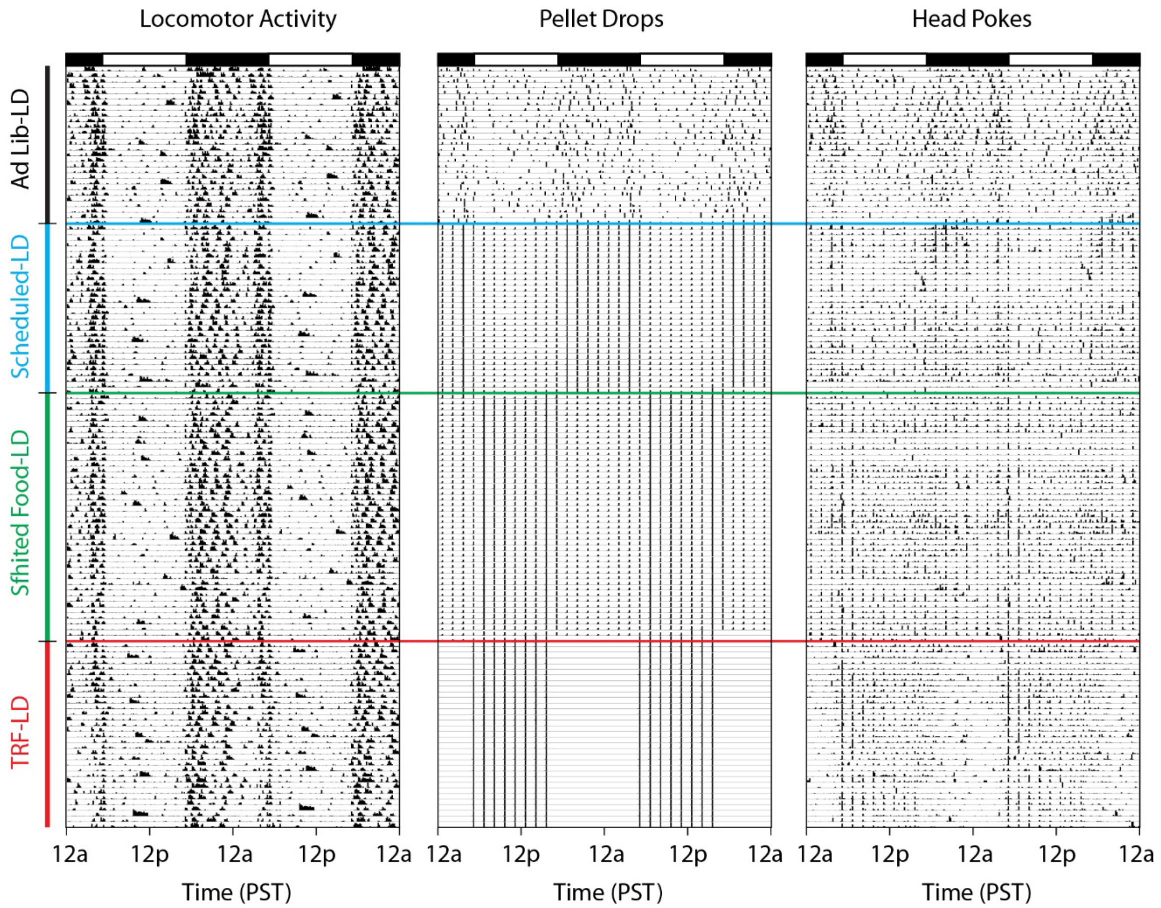

**Supplementary Figure S8.** Nocturnality index (median and 95% confidence interval) calculated as the ratio of activity during the dark to activity during the light. Mice were at least twice as active in the dark in all feeding conditions. In pairwise comparisons, groups with different letters are significantly different.

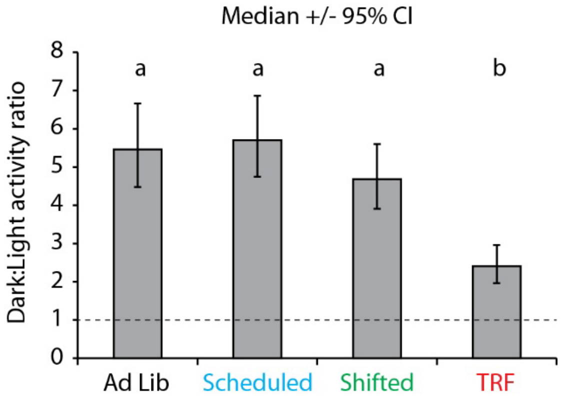


**Supplementary Figure S9.** Body weight (mean, SE) during Experiment 2. Triangles indicate PER2::LUC imaging days. Food schedule is indicated at the top (Ad Lib, Scheduled, TRF). DD began around week 8.

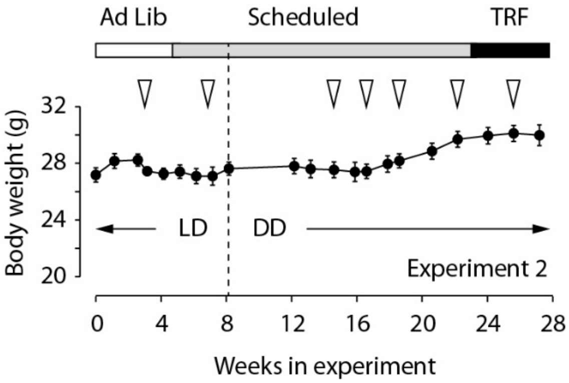


**Supplementary Figure S10.** In Experiment 2, the switch from Ad Libitum to Scheduled food in LD had small (~1h) effects on phase. These were not significant (paired t-test, left to right, $t_{11}$ = 1.89, 2.07, 1.67, and 0.54).

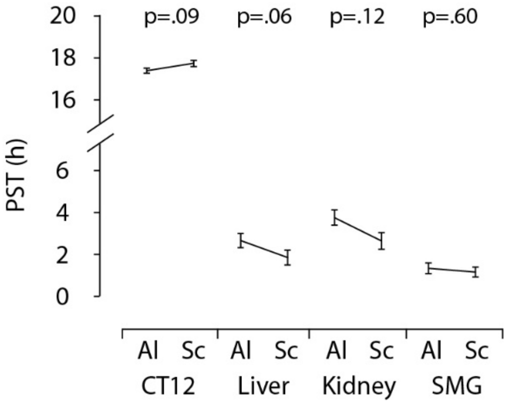

**Supplementary Figure S11.** Individual phases were calculated for each tissue and each mouse on each of the four measurement days in constant conditions. These phases were plotted either by food time (top) or by circadian time (bottom). When plotted by circadian time, vectors tended to be longer and more closely clustered, indicating that peripheral organs are entrained to the internal circadian system. In contrast, phases were widely distributed around the clock when analyzed by food time. Vectors show the resultant phase for each of the four imaging time points; vectors longer than the dotted circle (r=0.5) indicate significant clustering in phase (Rayleigh test, p<.05). P values next to each plot are the result of the Mardia-Watson-Wheeler circular test for differences in phase between imaging points ($\chi^2$ (df=6) for food time: kidney = 25.1, liver = 14.9, SMG = 25.5; for circadian time: kidney = 5.7, liver = 7.3, SMG = 10.7). Significant differences indicate no systematic clustering across the four measurement days.

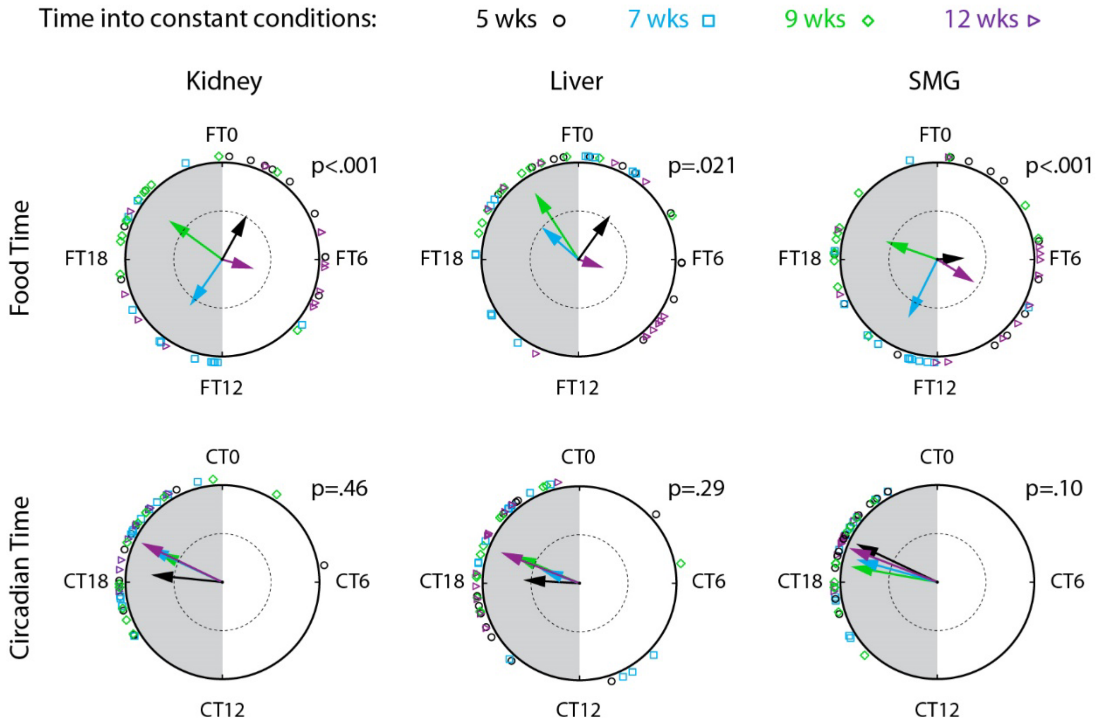

**Supplementary Figure S12.** Individual examples of peripheral clock phase against food schedule and individual rest-activity cycle show variability in the pattern of entrainment. Free-running activity patterns of two mice in constant conditions are shown in A and B. The food schedule is shown at the top. With intermittent assessment of peripheral clock phase, it is not possible to determine if changes in phase are due to advances and delays. The most parsimonious phase tracks are shown on the left. While the kidney and SMG both track with activity, the liver tracks with food (top) or with activity (bottom). The parsimonious tracks are determined by comparing two different tracks for each organ (C, D), one in which phase is assumed to follow food and one in which phase is assumed to follow activity. Insets show the mean resultant phase statistics (r, n=4/tissue) based on food phase or circadian phase for each mouse (see Fig. 6 legend for details).

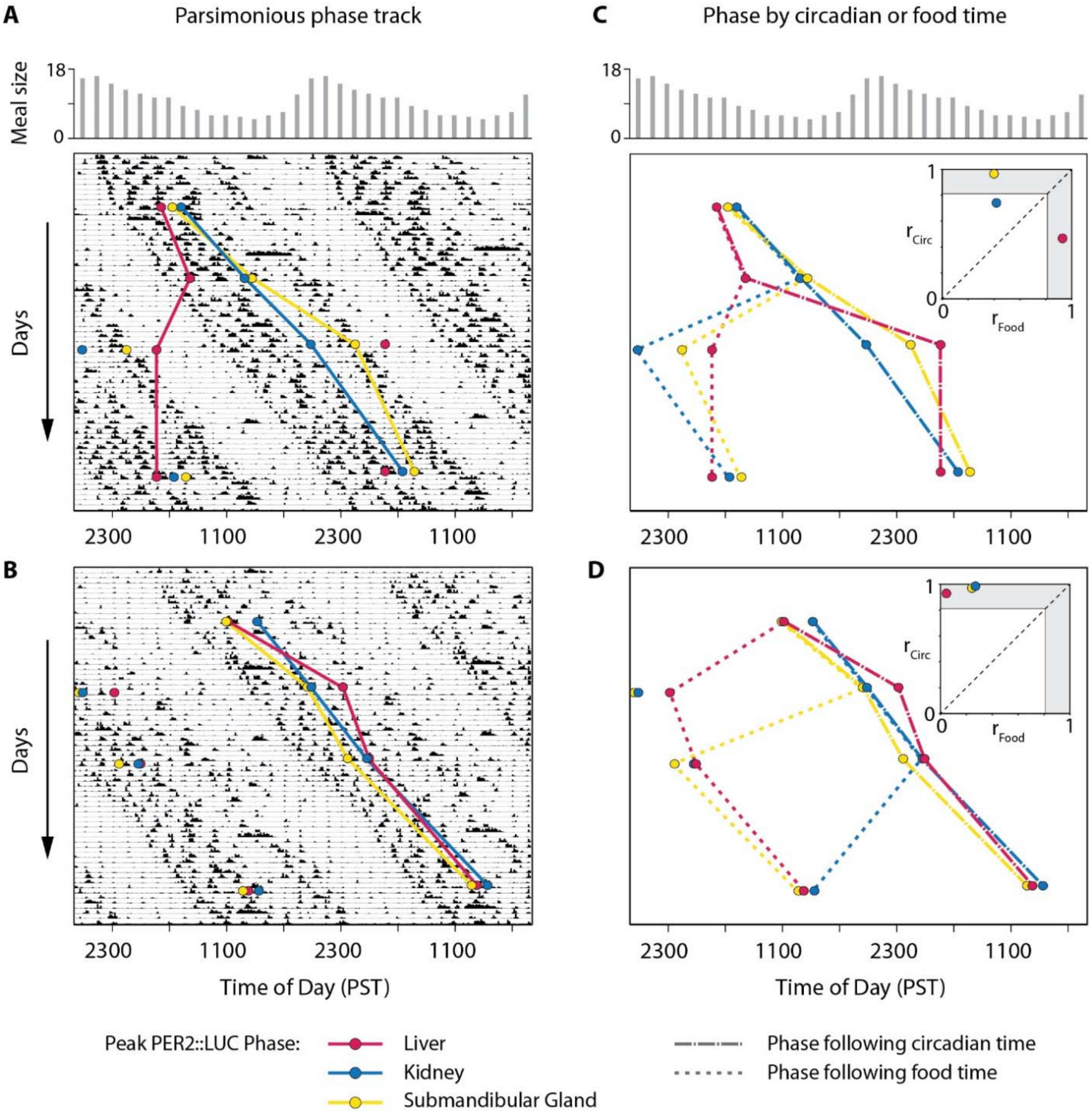

**Supplementary Figure S13.** Individual variability in response of locomotor activity rhythms to food schedules in constant dim red light. Simulated natural food schedules were not sufficient to entrain mice during constant conditions and all mice were free-running. When mice were then transferred to TRF with 12h food availability, circadian period significantly shortened (A, $t_{11}$ = 2.36, p=.038). Mice fell into three general categories: no change or longer period (B, n=3), shortened period (C, n=7), and entrained (D, n=2). In the actograms, grey shading indicates the food schedule. Peripheral organ phase is also shown. Note that PER2::LUC bioluminescence in the SMG always peaks in the middle of the animal's active phase. The liver and kidney transition from peaking during the active phase at the top to phase-locked to the end of food availability during TRF.

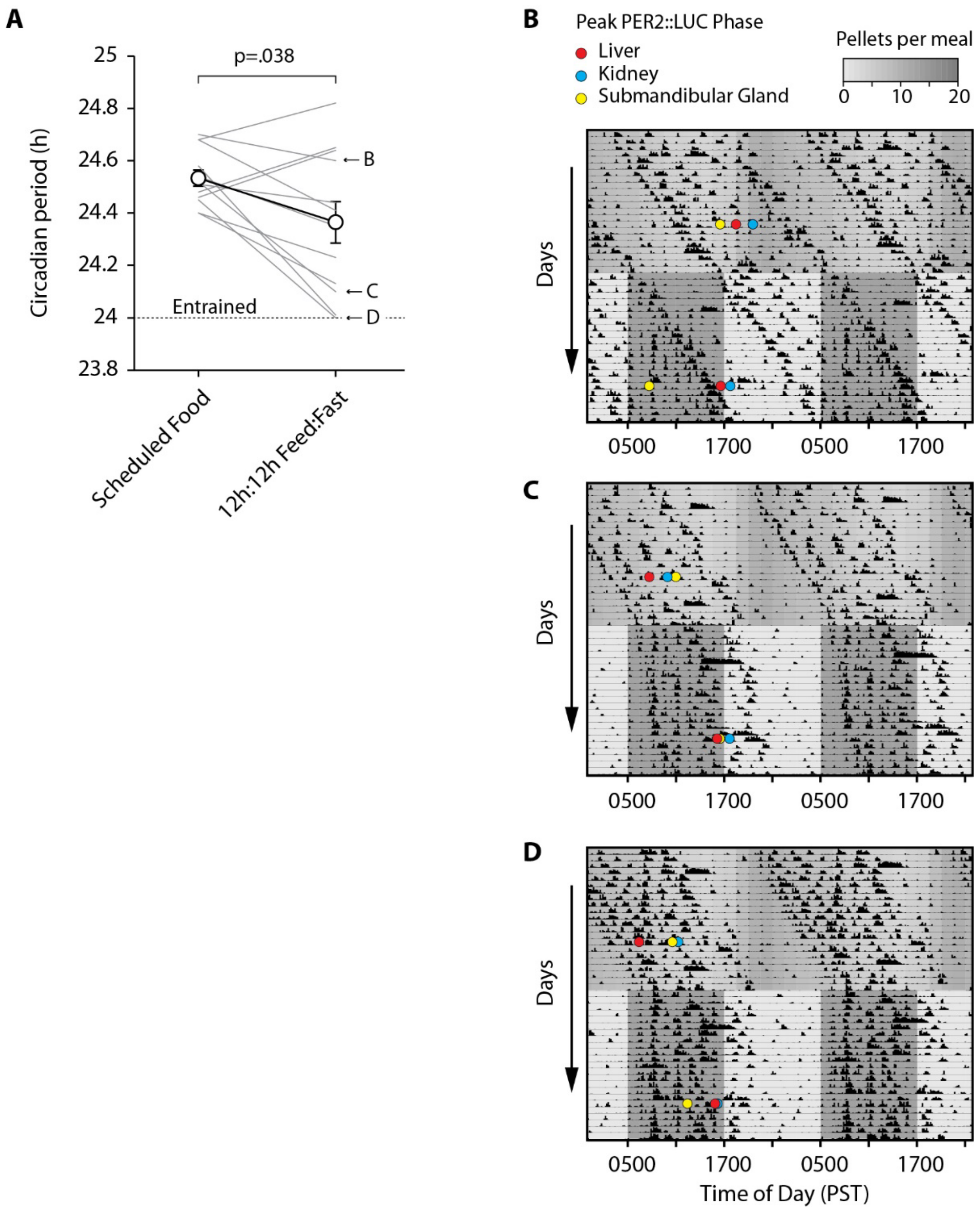

**Supplementary Figure S14.** Individual tissue phases and activity phase (CT12), in food time (A) or circadian time (B), in free-running mice exposed to TRF in constant dark conditions (food available from FT12 to FT0). Liver and kidney phases show tighter clustering with respect to food, while submandibular gland phase is more clustered with respect to circadian time. Circadian time is defined by activity, so no statistical test was performed for this. The vertical lines show the mean phase of each parameter during baseline LD Ad Lib conditions assuming that CT12 = FT12 = ZT12.

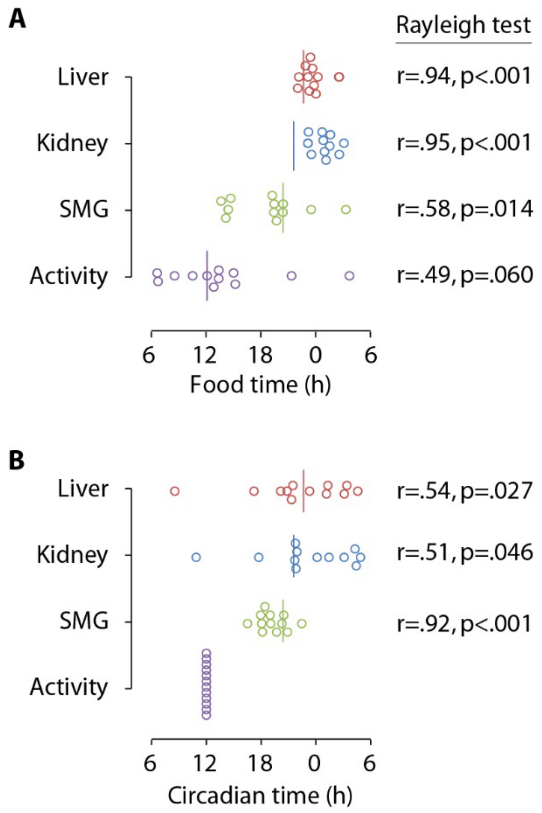

### Supplementary Tables

**Table S1**. In experiment 2, normalized PER2::LUC bioluminescence across the four tests (weeks 6-14 in DD) was analyzed by cosinor regression with independent variables of circadian time (CT), food time (FT), or both (12 mice × 4 tests × 6 bioluminescence measures = 288 points per organ). Mouse number was included as a random factor to account for repeated measures. The table shows goodness of fit and coefficient estimates for the different models. Regressions against CT outperformed regressions against FT as indicated by larger $R^2$ and smaller AICc. Though the effect of adding FT to the model was small, it significantly improved the model fit (Partial F test based on reduced SSE, Liver, $F_{8,268}$ = 6.8, p<.001; Kidney, $F_{8,268}$ = 4.9, p<.001; SMG, $F_{8,268}$ = 3.7, p<.001). Including time into constant darkness as a fixed factor did not affect $R^2$ because mean bioluminescence was normalized to 100% within mouse and within experimental day (6 measures per mouse).

| | Fixed Effects Included in Model | | |
|---|---|---|---|
| | CT | FT | CT+FT |
| Liver | $R^2$ = 0.36<br>AICc = 2869<br>cos(CT) $F_{1,285}$ = 12.1, p<.001<br>sin(CT) $F_{1,285}$ = 148, p<.001 | $R^2$ = 0.09<br>AICc = 2969<br>cos(FT) $F_{1,285}$ = 28.0, p<.001<br>sin(FT) $F_{1,285}$ = 0.7, p=.40 | $R^2$ = 0.47<br>AICc = 2810<br>cos(CT) $F_{1,283}$ = 19.0, p<.001<br>sin(CT) $F_{1,283}$ = 186, p<.001<br>cos(FT) $F_{1,283}$ = 56.5, p<.001<br>sin(FT) $F_{1,283}$ = 4.4, p=.037 |
| Kidney | $R^2$ = 0.52<br>AICc = 2767<br>cos(CT) $F_{1,285}$ = 22.3, p<.001<br>sin(CT) $F_{1,285}$ = 284, p<.001 | $R^2$ = 0.02<br>AICc = 2970<br>cos(FT) $F_{1,285}$ = 0.8, p=.38<br>sin(FT) $F_{1,285}$ = 4.8, p=.029 | $R^2$ = 0.56<br>AICc = 2740<br>cos(CT) $F_{1,283}$ = 25.4, p<.001<br>sin(CT) $F_{1,283}$ = 318, p<.001<br>cos(FT) $F_{1,283}$ = 4.2, p=.041<br>sin(FT) $F_{1,283}$ = 20.8, p<.001 |
| SMG | $R^2$ = 0.70<br>AICc = 2900<br>cos(CT) $F_{1,285}$ = 45.2, p<.001<br>sin(CT) $F_{1,285}$ = 621, p<.001 | $R^2$ = 0.04<br>AICc = 3232<br>cos(FT) $F_{1,285}$ = 10.4, p=.001<br>sin(FT) $F_{1,285}$ = 1.9, p=.17 | $R^2$ = 0.74<br>AICc = 2852<br>cos(CT) $F_{1,283}$ = 46.9, p<.001<br>sin(CT) $F_{1,283}$ = 730, p<.001<br>cos(FT) $F_{1,283}$ = 26.0, p<.001<br>sin(FT) $F_{1,283}$ = 21.6, p<.001 |

**Table S2**. Total number of fasting intervals (no pellets dropped) of different durations during Ad Lib baseline conditions in both experiments. The number of fasts occurring in the light phase are also shown.

Experiment 1 (8 d baseline in 10 mice = 80 mouse-days)

| | Number of fasting intervals | | | | | |
|---|---|---|---|---|---|---|
| | Total | >1h | >1.5h | >2h | >3h | >4h |
| Number | 1862 | 432 | 192 | 88 | 33 | 13 |
| Percentage | 100.0 | 23.2 | 10.3 | 4.7 | 1.8 | 0.7 |
| Number in light phase | 737 | 142 | 134 | 59 | 21 | 12 |
| Percentage | 39.6 | 7.6 | 7.2 | 3.2 | 1.1 | 0.6 |

Experiment 2 (10 d baseline in 12 mice = 120 mouse-days)

| | Number of fasting intervals | | | | | |
|---|---|---|---|---|---|---|
| | Total | >1h | >1.5h | >2h | >3h | >4h |
| Number | 2329 | 672 | 371 | 226 | 109 | 53 |
| Percent of total | 100.0 | 28.9 | 15.9 | 9.7 | 4.7 | 2.3 |
| Number in light phase | 951 | 408 | 226 | 132 | 57 | 35 |
| Percent of total | 40.8 | 17.5 | 9.7 | 5.7 | 2.4 | 1.5 |

**Table S3**. The number of long (>4h) fasts experienced by each mouse. Note that 20/22 mice had a >4h fast on half or fewer of their baseline days.

| | | Number of fasts > 4h in light phase | | | | | | | | | |
|---|---|---|---|---|---|---|---|---|---|---|---|
| | | 0 | 1 | 2 | 3 | 4 | 5 | 6 | 7 | 8 | 9 |
| Expt. 1 | # of mice | 3 | 5 | | 1 | 1 | | | | | |
| | Fasts/day | 0 | 0.125 | | 0.375 | 0.5 | | | | | |
| Expt. 2 | # of mice | 2 | 3 | 2 | 1 | 2 | | | | 1 | 1 |
| | Fasts/day | 0 | 0.1 | 0.2 | 0.3 | 0.4 | | | | 0.8 | 0.9 |
